# Open source timed pressure control hardware and software for delivery of air mediated distensions in animal models

**DOI:** 10.1101/2021.03.15.435466

**Authors:** Trishna Patel, Jamie Hendren, Nathan Lee, Aaron D Mickle

**Affiliations:** Department of Physiological Sciences, College of Veterinary Medicine, University of Florida, Gainesville, FL, United States; Department of Biomedical Engineering, Herbert Wertheim College of Engineering, University of Florida, Gainesville, FL, United States; Deparment of Neuroscience, College of Medicine, University Florida, Gainesville, FL, United States

**Keywords:** Visceromotor response, visceral pain, Arduino, solenoid

## Abstract

**Abstract:** Studying the visceral sensory component of peripheral nervous systems can be challenging due to limited options for consistent and controlled stimulation. One method for mechanical stimulation of hollow organs, including colon and bladder, are controlled distensions mediated by compressed air. For example, distension of the bladder can be used as an assay for bladder nociception. Bladder distension causes a corresponding increase in abdominal electromyography, which increases with distension pressure and is attenuated with analgesics. However, the hardware used to control these distensions are primarily all one-off custom builds, without clear directions how to build your own. This has made it difficult for these methods to be fully utilized and replicated as not everyone has access, knowledge and resources required to build this controller. Here we show an open-source Arduino based system for controlling a solenoid valve to deliver timed pressure distensions in the experimental model. This device can be controlled by one of two methods through direct TTL pulses from the experimenters data acquisition software (ex. CED Spike2) or by a graphical user interface, where the user can set the time before, during, and after distension as well as the number of cycles. This systems low cost and relative ease to build will allow more groups to utilize timed pressure distensions in their experiments.

*Specifications table:* 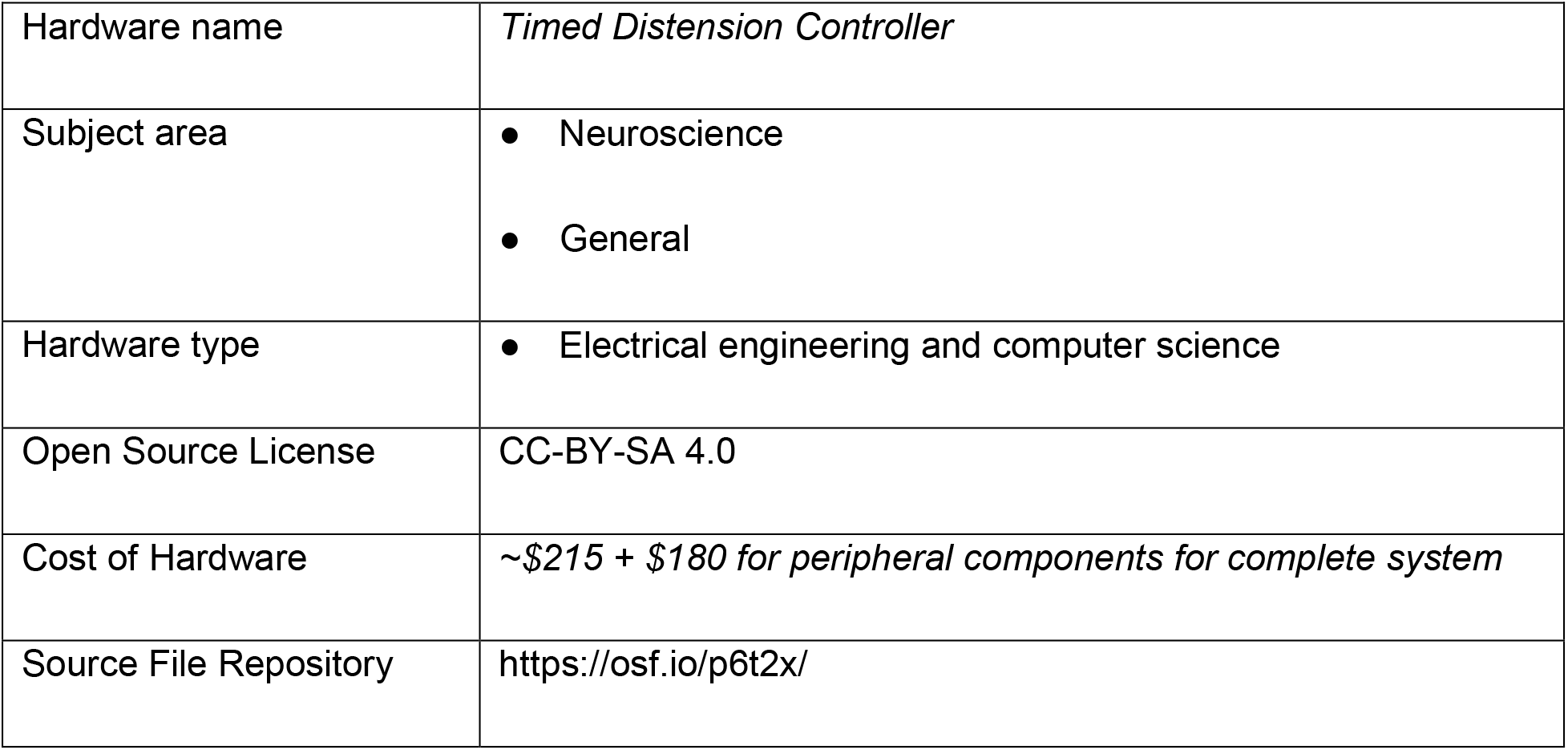 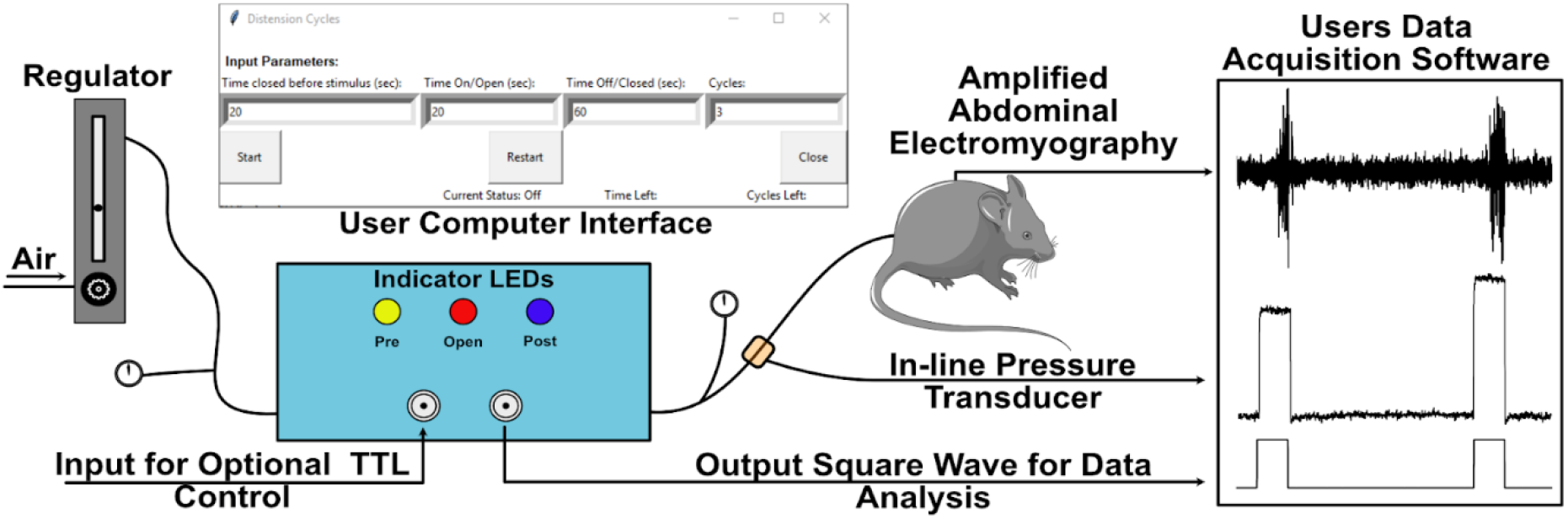

## 1. Hardware in Context

Normally we have very little perception of our bladder, intestine, and stomach as they function to extract nutrients and remove the body’s waste. However, for greater than 20% of the world’s population discomfort and/or pain accompany these normal physiological activities [1]. Unfortunately, the mechanisms surrounding these diseases, like irritable bowel syndrome, interstitial cystitis, and bladder pain syndrome have yet to be elucidated. One tool neuroscientists use to assess the sensory function of visceral structures is hollow organ distension which is a validated and widely used technique in both animal models [2–5] and human studies [6–8]. Graded distension directly with air or via a balloon can provide organ specific stimuli that is critical for studying these structures under normal and disease conditions.

These experimental distensions are typically controlled with either by manual opening of a valve or via an electronic control system [7, 9]. Using an electronic control system has a number of advantages including consistent timing of distention series, almost immediate on and off times, as well as alignment with other stimuli or experimental interventions. However, the use of these control systems has been limited as the devices described in the literature are all custom built with no published designs and usually custom made by various university machine shops. The exception is the timed pressure controller published by Andersen, Ness and Gebhart (1987) [9]. To build this system requires expertise and skill with electronics that is normally not found in neuroscience or biology labs. Outsourcing the build increases the cost and thus limits the number of groups who can build their own controller. With the increased availability and decreased cost of off the shelf microcontroller boards, like the Arduino Uno, there are now simpler ways of building and using a time distension controller.

Here we describe an easy to build, low-cost timed distension control device and peripheral components needed to fully use the system to study visceral sensory systems on rodents. This system can be assembled with limited tools and technical knowledge. This system will make it easier to utilize controlled distension stimulation in labs that study visceral organ physiology. We have designed two different ways to operate the Arduino controlled solenoid. The first is a standalone user-interface that uses the open-source programing language Python to control distention (variables duration, time between distensions, and number of trials). The second mode of operation uses transistor-transistor logic (TTL) to control the solenoid with other lab software/hardware, which allows precise coordination of distension with commonly used lab equipment and other experimental interventions.

## 2. Hardware description

Our timed distension controller uses an Arduino microcontroller to control a relay module which opens and closes a solenoid valve. The electronics and solenoid are contained within a 3D printed case with removable lid (Figure 1a and b). Opening the solenoid allows pressurized air to flow to the target/animal and distend the hollow organ or balloon (Figure 1c). The pressure of distension is controlled by a flowmeter regulator and is changed manually for different distension pressures (Figure 1c). The timing of the distension can be controlled using the python-based user interface (Figure 1d) or if precise coordination with other software/stimuli is needed, it can be triggered using a TTL pulse. Three LEDs on the outside of the case illuminate during each stage of the distension; yellow in the pre-distension time period, red during distension, and blue during the post-distension time.

**Figure 1.**
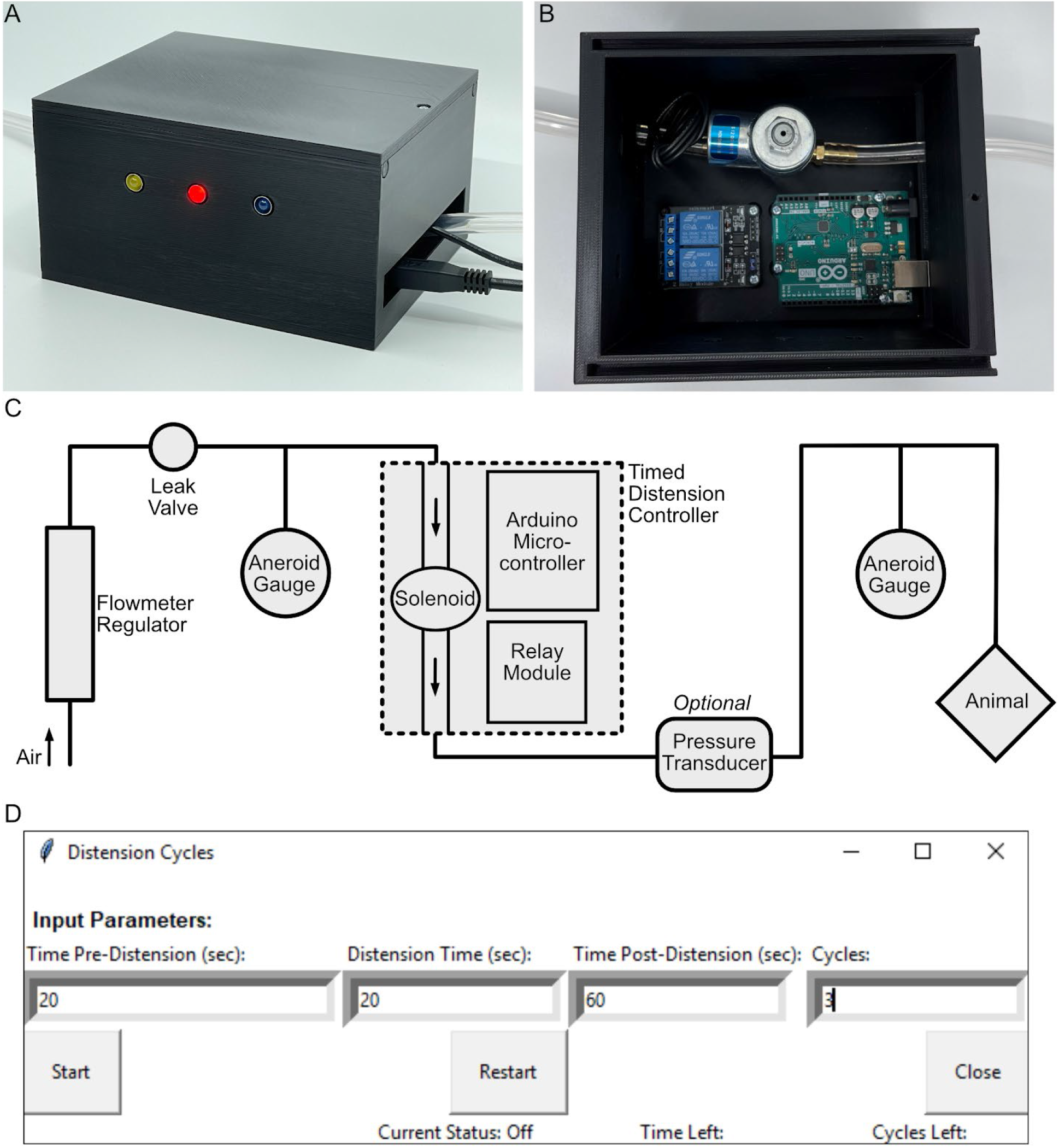
Timed Distension Controller and User Interface. The timed distension controller uses an Arduino microcontroller to control a relay module which opens and closes a solenoid valve (**A and B**). We have designed a 3D printable box to mount the electronics and solenoid. There are three LEDs that indicate what distension phase the device is in, closed before valve opening (yellow), time open (red) and time closed (blue). (**C**) This experimental diagram demonstrates how the timed distension controller fits within the entire distension workflow from compressed air tank to animal. (**D**) A screenshot of the user interface show the input parameters, time closed before stimulus (pre-distension time), time open (stimulus time), time off (post-distension time) and cycles (the number of times it is repeated).

The primary advantages of using the system described here over other systems are the availability, cost, and ability to control distention with a TTL interface. The only other published system is more than 30 years old and would be challenging for a non-electrical engineer to reproduce [9]. Newer systems have been used but the designs have not been published [10, 11]. If you can find someone to build one of these previously used systems, the cost is significantly higher than this system, with quotes in the $2000 to $3000 range. This cost primarily due to have to paying an engineer someone to build and potentially redesign the controller. In some previously described timed distension control devices the stimulus is timed with direct inputs [10, 11] and in others it can be triggered by TTL input [12–14]. For this device we have the option of either direct input control or triggering valve opening with TTL pulses for controlling distension with other software and/or to coordinate with other experimental interventions.

- This device is easy to build and with basic soldering skills any lab should be able to build their own.
- This device is cost effective with all the components costing around ~$200 compared to ~$2000-3000 obtained quotes to build something similar by a university machine shop core, where the plans are not publicly available.
- The design and components are easily accessible and can be modified to fit specific experimental needs.
- Operation modes are flexible for multiple applications with or without additional equipment.

## 3. Design files

### 3.1 Design Files Summary

| Design file name | File type | Open source license | Location of the file |
| --- | --- | --- | --- |
| Wiring Diagram | .PDF | CC-BY-SA 4.0 | https://osf.io/p6t2x/ |
| TPC-Case | .STEP | CC-BY-SA 4.0 | https://osf.io/p6t2x/ |
| TPC-Case | .STL | CC-BY-SA 4.0 | https://osf.io/p6t2x/ |
| TPC-Lid | .STEP | CC-BY-SA 4.0 | https://osf.io/p6t2x/ |
| TPC-Lid | .STL | CC-BY-SA 4.0 | https://osf.io/p6t2x/ |
| TCP_GUIv1.2 | .PY | CC-BY-SA 4.0 | https://osf.io/p6t2x/ |
| TPC_TTL_Triggering | .INO | CC-BY-SA 4.0 | https://osf.io/p6t2x/ |
| Air Flow Diagram | .PDF | CC-BY-SA 4.0 | https://osf.io/p6t2x/ |

### 3.2 Description of each file

- Wiring Diagram – PDF file with a wiring diagram to aid in assembly of the device.
- 3D Case and Lid Files – The STL files for 3D printing the timed distension controller case. The CAD file (.STEP format) has also been included if you want to make modifications.
- TCP_GUI_v1.2– Python code for the user interface to control the timed pressure controller.
- TCP_TTL_Triggering – Arduino code for the timed pressure controller TTL mode of operation.

## 4. Bill of Materials

*Please see attached editable spreadsheet file with links.*

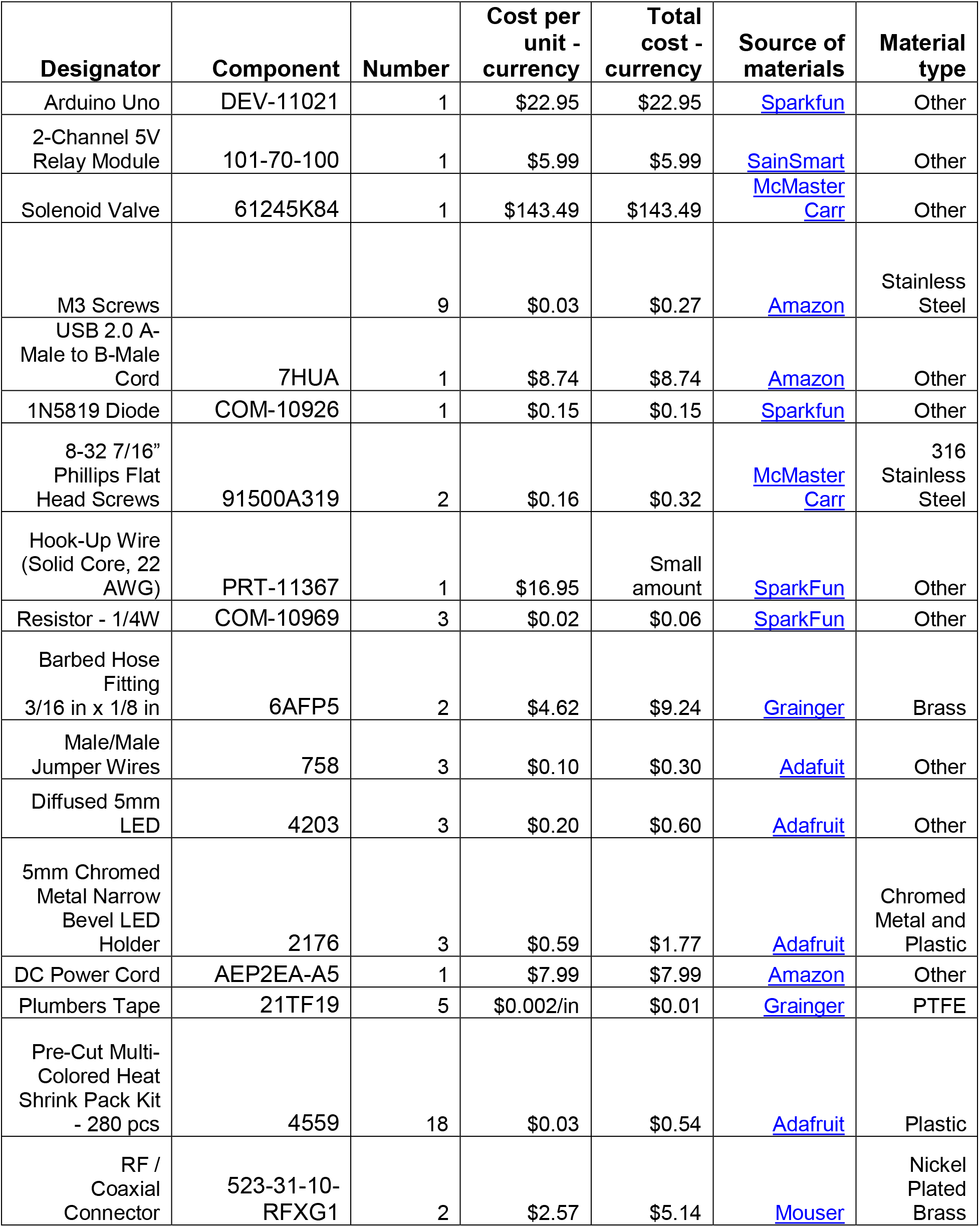
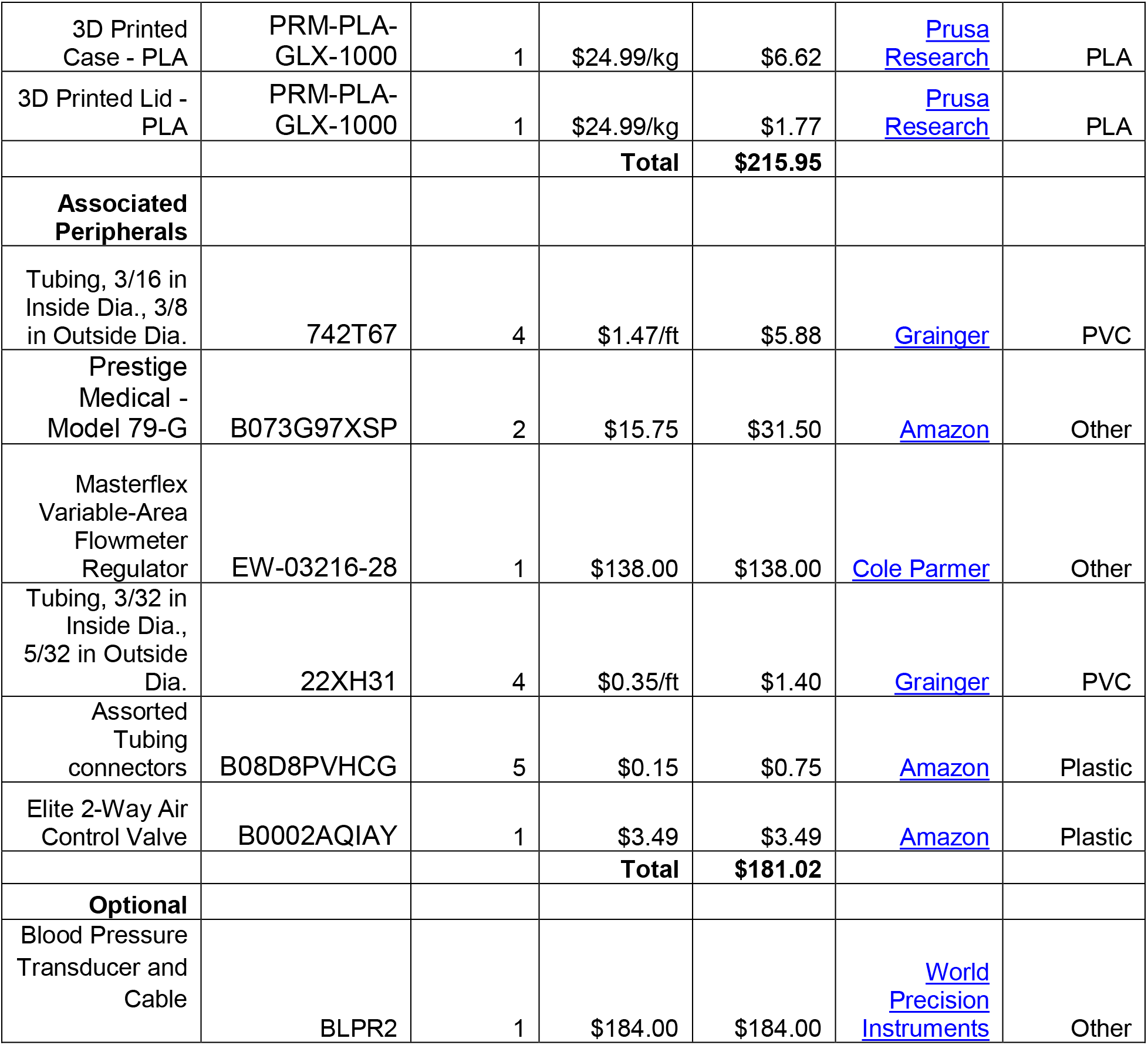

| Designator | Component | Number | Cost per unit - currency | Total cost - currency | Source of materials | Material type |
| --- | --- | --- | --- | --- | --- | --- |
| Arduino Uno | DEV-11021 | 1 | \$22.95 | \$22.95 | Sparkfun | Other |
| 2-Channel 5V Relay Module | 101-70-100 | 1 | \$5.99 | \$5.99 | SainSmart | Other |
| Solenoid Valve | 61245K84 | 1 | \$143.49 | \$143.49 | McMaster Carr | Other |
| M3 Screws | | 9 | \$0.03 | \$0.27 | Amazon | Stainless Steel |
| USB 2.0 A-Male to B-Male Cord | 7HUA | 1 | \$8.74 | \$8.74 | Amazon | Other |
| 1N5819 Diode | COM-10926 | 1 | \$0.15 | \$0.15 | Sparkfun | Other |
| 8-32 7/16" Phillips Flat Head Screws | 91500A319 | 2 | \$0.16 | \$0.32 | McMaster Carr | 316 Stainless Steel |
| Hook-Up Wire (Solid Core, 22 AWG) | PRT-11367 | 1 | \$16.95 | Small amount | SparkFun | Other |
| Resistor - 1/4W | COM-10969 | 3 | \$0.02 | \$0.06 | SparkFun | Other |
| Barbed Hose Fitting 3/16 in x 1/8 in | 6AFP5 | 2 | \$4.62 | \$9.24 | Grainger | Brass |
| Male/Male Jumper Wires | 758 | 3 | \$0.10 | \$0.30 | Adafruit | Other |
| Diffused 5mm LED | 4203 | 3 | \$0.20 | \$0.60 | Adafruit | Other |
| 5mm Chromed Metal Narrow Bevel LED Holder | 2176 | 3 | \$0.59 | \$1.77 | Adafruit | Chromed Metal and Plastic |
| DC Power Cord | AEP2EA-A5 | 1 | \$7.99 | \$7.99 | Amazon | Other |
| Plumbers Tape | 21TF19 | 5 | \$0.002/in | \$0.01 | Grainger | PTFE |
| Pre-Cut Multi-Colored Heat Shrink Pack Kit - 280 pcs | 4559 | 18 | \$0.03 | \$0.54 | Adafruit | Plastic |
| RF / Coaxial Connector | 523-31-10-RFXG1 | 2 | \$2.57 | \$5.14 | Mouser | Nickel Plated Brass |
| 3D Printed Case - PLA | PRM-PLA-GLX-1000 | 1 | \$24.99/kg | \$6.62 | Prusa Research | PLA |
| 3D Printed Lid - PLA | PRM-PLA-GLX-1000 | 1 | \$24.99/kg | \$1.77 | Prusa Research | PLA |
| | | | **Total** | **\$215.95** | | |
| **Associated Peripherals** |  |  |  |  |  |  |
| Tubing, 3/16 in Inside Dia., 3/8 in Outside Dia. | 742T67 | 4 | \$1.47/ft | \$5.88 | Grainger | PVC |
| Prestige Medical - Model 79-G | B073G97XSP | 2 | \$15.75 | \$31.50 | Amazon | Other |
| Masterflex Variable-Area Flowmeter Regulator | EW-03216-28 | 1 | \$138.00 | \$138.00 | Cole Parmer | Other |
| Tubing, 3/32 in Inside Dia., 5/32 in Outside Dia. | 22XH31 | 4 | \$0.35/ft | \$1.40 | Grainger | PVC |
| Assorted Tubing connectors | B08D8PVHCG | 5 | \$0.15 | \$0.75 | Amazon | Plastic |
| Elite 2-Way Air Control Valve | B0002AQIAY | 1 | \$3.49 | \$3.49 | Amazon | Plastic |
| | | | **Total** | **\$181.02** | | |
| **Optional** |  |  |  |  |  |  |
| Blood Pressure Transducer and Cable | BLPR2 | 1 | \$184.00 | \$184.00 | World Precision Instruments | Other |

### Additional Notes on Component Selection

The electrical air directional control solenoid valve was chosen because it operates on 12V, is normally closed, and has a vent to release air pressure from the OUT tubing allowing the distension to quickly end after the value closes. When we originally purchased the valve, it was cheaper than the listed price on the bill of materials. There may be other more cost-effective options now available.

*Tools needed that are not included on bill of materials list:*

- Soldering iron, Teflon tape, crescent wrench, screwdrivers, scissors, wire strippers, and head gun.
- 3D printer - Parts could be ordered from a commercial 3D part printer

- All parts were printed on a Prusa MK3s, STL file was prepared for printing using Prusa Slicer 2.3, the standard 0.30 mm Draft setting was used (with a layer height of 0.3 mm, an infill density of 20%, a print temperature of 210 °C, and a bed temperature of 60 °C).

## 5. Build Instructions

### 5.1 Assembly of timed distension controller hardware

All circuit components are detailed in the bill of materials. Use the wiring diagram to aid in wiring and assembly of the timed distension controller. 3D-print the case and lid prior to starting assembly.

1. Attach Arduino Uno and relay module to case mounts using 6 mm M3 screws (Figure 2a).
2. Prepare Solenoid before mounting in case (Figure 2b).

a. Wrap both barbed hose fittings in plumber’s Teflon tape.
b. Screw in each barbed hose fitting onto solenoid outlets.
c. Attach 3/16 inch inner diameter tubing onto the barbed portion of each fitting.
3. Mount solenoid to case.

a. Feed tubes through case openings. The tube on the wiring side should go through the round opening and the other tube through the rectangular opening. There is also a flow arrow on the solenoid. The in tube should enter through the smaller hole on the case and the exit tube will go out the larger rectangular hole. It is important to mount the solenoid in the correct direction as the OUT side of the value has a value to release air when the solenoid is closed. This will dissipate distension when the stimulus is removed.
b. Line up solenoid over mounting holes.
c. Holding solenoid in place with one hand, flip case over and insert an 8-32 screw into each hole, and tighten the screws to attach solenoid.
4. Insert the LED holders, threaded side first, into each of the three LED holes (three uniform holes on the long side of the case). Secure the LED holder in its designated hole with its washer.
5. Wire together Arduino and relay module using male-to-male jumper wires.

a. Connect relay GND to Arduino GND, relay IN1 to Arduino 13, and relay VCC to Arduino 5V (Figure 2c).
b. Use a jumper to connect VSS to JD-VCC on the relay module to power the relay with the Arduino.
6. Prepare RF/coaxial connector for mounting (Figure 2d).

a. Cut one 3-inch segment and one 5-inch segment of solid core wire and strip ~ 1 cm off each end of both wires. Choose two different colors to aid in identifying the different pin outputs after RF connector is prepared.
b. Solder the 3-inch wire to ground (flat) pin on connector and the 5-inch wire to the positive (round-center) pin. Black wire was used for ground and red for positive in the images.
c. Cover soldered connection and any exposed portion of the pins with heat shrink, using the smallest size that will fit over the solder. Apply hot air to secure the heat shrink. Heat shrink is used to prevent any exposed metal/wire/solder from touching and to support any brittle soldering connections.
d. Repeat steps 6a-c. with second RF connector.
7. Mount and install RF/coaxial connectors.

a. Insert RF/coaxial connectors into the two slots on the short side of the case, feeding the wires in first. Secure using associated washer.
b. Insert positive wire for the right RF/coaxial connector into Arduino pin 2 and the positive wire for the left RF connector into Arduino pin A0.
c. Twist the ground wires for the RF/coaxial connector together. Cut another 3-inch segment of solid core wire and strip ~ 1 cm from each end. Twist this 3^rd^ wire around the other 2 (Figure 2e). Solder this connection together, and cover with heat shrink, applying hot air to secure. The end of this wire should then be connected to an Arduino GND pin (Figure 2f).
8. Prepare red, yellow, and blue LEDs for mounting (Figure 2g).

a. For one LED, feed LED legs through plastic portion of LED holder.
b. Bend ground (short) leg of LED, cut a 3-inch segment of solid core wire and strip ~ 1 cm from each end. Twist one end of the wire with the LED ground leg.
c. Solder that connection and cover with heat shrink, applying hot air to secure.
d. Repeat steps 8 a-c. with other two LEDs.
e. For each LED, connect positive (long) leg to its respective resistor by twisting the LED’s positive leg and one of the resistor’s legs together and soldering that connection. The LEDs in our circuit require resistors to limit current flow to each LED. This ensures that the LEDs will not burn out. The red and yellow LEDs have similar voltage requirements and need one 220Ω resistor each. The blue LED has a higher voltage requirement and will need one 100Ω resistor. Cover soldered connections with heat shrink and secure using hot air.
f. Next, gather the three ground wires from each LED and twist them together. Cut a 3-inch segment of solid core wire and strip ~1 cm off each end, then twist one end of this wire together with the other three wires. Solder this connection and cover with heat shrink, applying hot air to secure.
g. Next, cut three 3-inch pieces of solid core wire, stripping ~ 1 cm off each of all three wires. For each LED, twist the free leg of their respective resistor together with one piece of solid core wire, solder the connection, cover with heat shrink, and apply hot air to secure.
9. Mount and install the LEDs.

a. Mount the LEDs into their holders in the case from the inside, arranged so that the yellow LED is on the left, red in the middle, and blue on the right.
b. Connect the LED’s ground wire to Arduino GND pin, the yellow LED’s positive wire to Arduino pin 6, the red LED’s positive leg to Arduino pin 8, and the blue LED’s positive leg to Arduino pin 10 (Figure 2h).
10. Prepare solenoid wires to be connected to relay unit.

a. Cut end off DC power cord plug, cut a slit from exposed end up ~4 inches, and pull out the ground and positive wires from inside the cord. Strip ~ 1 cm from the ends of both positive and negative wires.
b. Place heat shrink on the ground cord for later use. Twist black ground wire together with the ground leg of a diode and solder the connection, making sure to keep the heat shrink away from the connection otherwise the heat shrink will pre- emptively shrink. The diode prevents current discharge from coming back through the circuit and damaging the circuit.
c. Feed power cord and attached diode inside case through large rectangular opening on the case’s front face, making sure to pull extra cord through to allow for mobility in the wire.
d. Pick one of the solenoid wires and place a piece of heat shrink on it. Then, wrap the wire around the soldered DC power ground-diode connection. (Figure 2i). Solder the solenoid wire into place. Push the two pieces of heat shrink, one on the solenoid wire and the other on the ground wire, together to cover as much of the soldered connection as possible. Cover the rest of the solder with heat shrink cut to diode leg size and slid over the diode. Apply hot air to shrink all the wrap.
e. Put ½ of a piece of heat shrink over the diode, pushed as far down the diode as possible, and another full piece down the second solenoid wire. Twist the positive leg of the diode together with the second solenoid wire. Then, cut a 3-inch piece of solid core wire, stripping ~ 1 cm from each end, and twist one of the wires ends together with the diode and solenoid wire. Solder this connection together, then push the two pieces of heat shrink together to cover as much solder as possible. Put a third piece of heat shrink over the solid core wire and push it down to cover the rest of the exposed solder. Apply hot air to shrink all the wrap (Figure 2j).
f. Bend the exposed metal of the power cord’s positive wire and the free end of the solid core wire at a 90-degree angle. Loosen the screws on the COM (middle) port and the NO (normally open; right) ports of the K1 channel. Insert the solid core wire into the COM port, tighten the port’s screw down, and tug gently to ensure it is securely connected. Insert the positive power wire into the NO (normally open) port, tighten the port’s screw down, and tug gently to ensure it is securely connected (Figure 2k).
11. Label the left (going to pin A0) RF/coaxial connector with “IN” and the right (going to pin 2) RF/coaxial connector with “OUT” (Figure 2l).
12. Slide case lid on and use a 16 mm M3 screw to secure lid into place.

**Figure 2.**
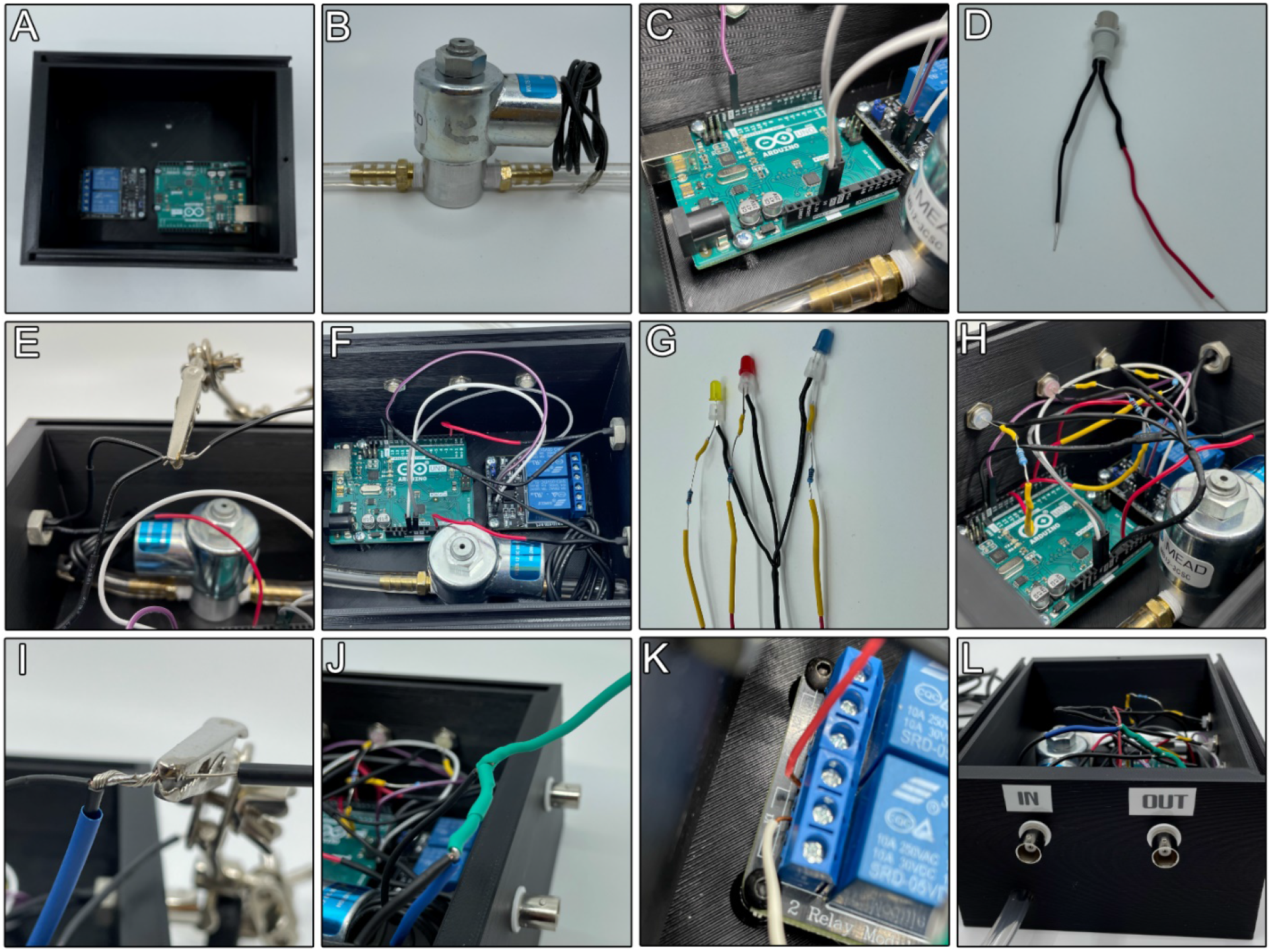
Assembly Photos of Timed Distension Controller Hardware. The timed distension controller hardware consists of an Arduino Uno microcontroller and a 2-channel relay that are aired together, and a solenoid mounted to the base of a 3D printed case (**A, B and C**). The hardware also includes mounted RF/coaxial connectors and LEDs, which are all wired to the Arduino (**D, E, F, G and H**). Lastly, there is a DC power and diode connected to the two solenoid wires, the output of which are then plugged into the relay module (**I, J and K**). To complete the hardware, labels are added to the outside of the box to distinguish the two RF/coaxial connectors based on their Arduino wiring (**L**).

### 5.2 Assembly of complete timed distension controller system with peripherals

1. The air flow diagram in Figure 1c shows how to connect the tubing from the compressed air source to the animal. There is an additional ‘air flow diagram’ file in the build files.
2. Connect the air cylinder regulator to flow meter regulator using appropriate tubing.
3. Then using 3/16 inch interior diameter tubing, connect the flow meter to the leak valve. This allows for a small amount of leak making it easier to adjust the pressures and limiting the pressure that can go to the animal.
4. From the leak valve using the 3/16 inch inner diameter tubing run to a T connector. One side of the T goes to the first aneroid gauge. The other side of the T goes in to the IN inlet of the solenoid valve in the time distension controller.
5. From the OUT of the solenoid valve run the 3/16 inch interior diameter tubing to the barbed 3/16 to 3/32 reducer.
6. From the other side of the reducer, now using the smaller 3/32 inch inside diameter tubing, run to a T connector where one side will go to the second android gauge and the other will go either to the optional pressure sensor or to the animal.
7. Coming out of the optional pressure sensor we run the tubing to the animal. The final adaptor will depend on what you are using to distend. In our case we used a barbed to slip Luer lock adaptor to connect to a catheter.

### 5.3 Arduino Software Installation

There are two modes of operation independent control, through graphical user interface (GUI)-based or TTL triggered software. These require different software to be loaded on the Arduino so either follow 5.3.1 or 5.3.2. The mode of operation can be changed later if needed.

#### 5.3.1 Installation of independent GUI-based control

1. Download the Arduino IDE desktop app from arduino.cc (https://www.arduino.cc/).

a. Follow the directions for download options based on computer specs.
2. Install and open the Arduino IDE.
3. Connect the Arduino hardware to the PC via USB cable.
4. Arduino Startup

a. Configure the board

i. Tools > Board
ii. Select which Arduino board is being used
b. Select appropriate Port

i. Tools > Port
ii. Select the port indicating the Arduino Uno
5. In the Arduino IDE, run “Standard Firmata”.

a. File > Examples > Firmata > Standard Firmata
b. Verify and Upload
c. This file must always be running for the python scripts to run,
6. Install Python

a. Go to python.org (https://www.python.org/downloads/)
b. Download the latest version (we are using version 3.8)
c. Open Software
7. In Python, install the pyFirmata package.

a. Go to pypi.org (https://pypi.org/project/pyFirmata/)
b. Copy “pip install pyFirmata”
c. Open Python and in the IPython console paste and run the code
d. Restart Python to use the added package
8. Upload the “TCP_GUIv1.2” to Python terminal.

a. Open code in python terminal, we used Spyder 2.8 IDE.
b. In line 23 of the code, change the port from ‘COM3’ to whichever port the user is working with, i.e. the port that was selected in the Arduino IDE.
9. Save and run the code (Section 6.1.1)

#### 5.3.2 Installation of TTL triggered operation

1. Download the Arduino IDE desktop app from arduino.cc (https://www.arduino.cc/).

a. Follow the directions for download options based on computer specs.
2. Install and open the Arduino IDE.
3. Connect the Arduino hardware to the PC via USB cable.
4. Arduino Startup

a. Configure the board

i. Tools > Board
ii. Select which Arduino board is being used
b. Select appropriate Port

i. Tools > Port
ii. Select the port indicating the Arduino Uno
5. In the Arduino IDE, run “TPC_TTL_Triggering

a. Verify and Upload

## 6. Operation Instructions

### 6.1.1 Operation of timed distension controller with of GUI -based control

1. After running the code (Step 5.3.1). A pop-up labeled “Distension Cycles” will appear (Figure 1d). This window contains the 4 different input variables, Time Pre-Distension, Distension Time, Time Post-Distension and Cycles.
2. When operating the program, the “Start” button will begin to run the software, and the “Time Left:” and “Cycles Left:” fields in the bottom left corner of the pop-up will auto populate to allow you to keep track of the hardware status and cycle information
3. The “Restart” button can be used if the cycle set needs to be stopped or reset. This will return the valve to the closed state.
4. When finished use the “Close” button to exit out of the software.

### 6.1.2 Operation of timed distension controller with of TTL triggering

1. After uploading the “TPC_TTL_Triggering” the hardware will be set up to receive TTL signals to open the solenoid.
2. Opening of the valve and distension can be triggered by a TTL pulse greater that 4V. This can be changed in the “TPC_TTL_Triggering file if your system delivers a different voltage level.
3. When a pulse (>4 V) is delivered the solenoid will open. Programing the TTL pulses will depend on the software and hardware being used. If you need a simple TTL triggering device to coordinate multiple TTL pulses we suggest the [Arduino TTL Pulse Generator and Controller](https://optogeneticsandneuralengineeringcore.gitlab.io/ONECoreSite/projects/Arduino%20TTL%20Pulse%20Generator%20and%20Controller/) from the Optogenetics and Neural Engineering Core at the University of Colorado Anschutz Medical Campus. This can coordinate multiple TTL pulses like we used in the validation (section 7).

### 6.2.1 Safety Considerations

There are no major safety concerns with the operation of this device. However, we do test the device every time we use it on a small balloon (glove finger) to make sure the valve is opening and closing when it should be. We do this because we don’t want to accidently distend the animal when not intending to, which can lead to organ damage or cause undo harm to the animal. The leak valve will also help reduce chances of accidently distending at high pressures. We recommend unplugging the solenoid and Arduino when not in use.

## 7. Validation and Characterization

In order to validate the timed pressure controller, we performed a bladder distension evoked visceral motor response assay [3, 10, 11, 15]. This experiment is an assay of bladder nociception in mice [3, 11]. Increasing bladder distension pressures result in a corresponding increase in electromyography (EMG) activity in the superior oblique abdominal muscles. This reflex is termed a visceral motor response. This visceromotor response can be attenuated with the use of analgesics suggesting it can be used as an indirect measure of bladder nociception in mice [3]. A detailed explanation and descriptive methods have been published in the Journal of Visualized Experiments [11].

Here we used wild-type C57/B6 mice to demonstrate the functionality of the timed distention controller and associated software to increases in pressure (20, 30, 40, 50, 60, 70 mmHg). After a transurethral catheter is inserted, EMG electrodes are placed in the superior oblique abdominal muscles and the correct anesthesia plane is maintained; we then proceeded with a distension protocol [11]. We started by adjusting the distension pressure using the flowmeter regulator and the pre-solenoid aneroid gauge to 20 mmHg. Once the pressure is set, we entered our input parameters in the user interface (Time closed before stimulus = 20, Time on/open = 20, Time off/closed = 60 and Cycles = 18) (Figure 1d). After clicking ‘start’ we observed the animal and the post-solenoid aneroid gauge and once the distension started we adjusted the pressure with the flowmeter regulator if necessary so that the post-solenoid aneroid gauge indicated 20 mmHg. After 3 cycles, we then increased by 10 mmHg and repeated the same process up to 70 mmHg. Using an amplifier (Grass P511), Micro 1404 digitizer (CED) and Spike2 (CED) software, and a pressure transducer, we recorded EMG responses from the superior oblique abdominal muscles, solenoid pressure output, and the coaxial voltage output from the timed pressure controller, which indicates that the valve is open (Figure 3a).

**Figure 3.**
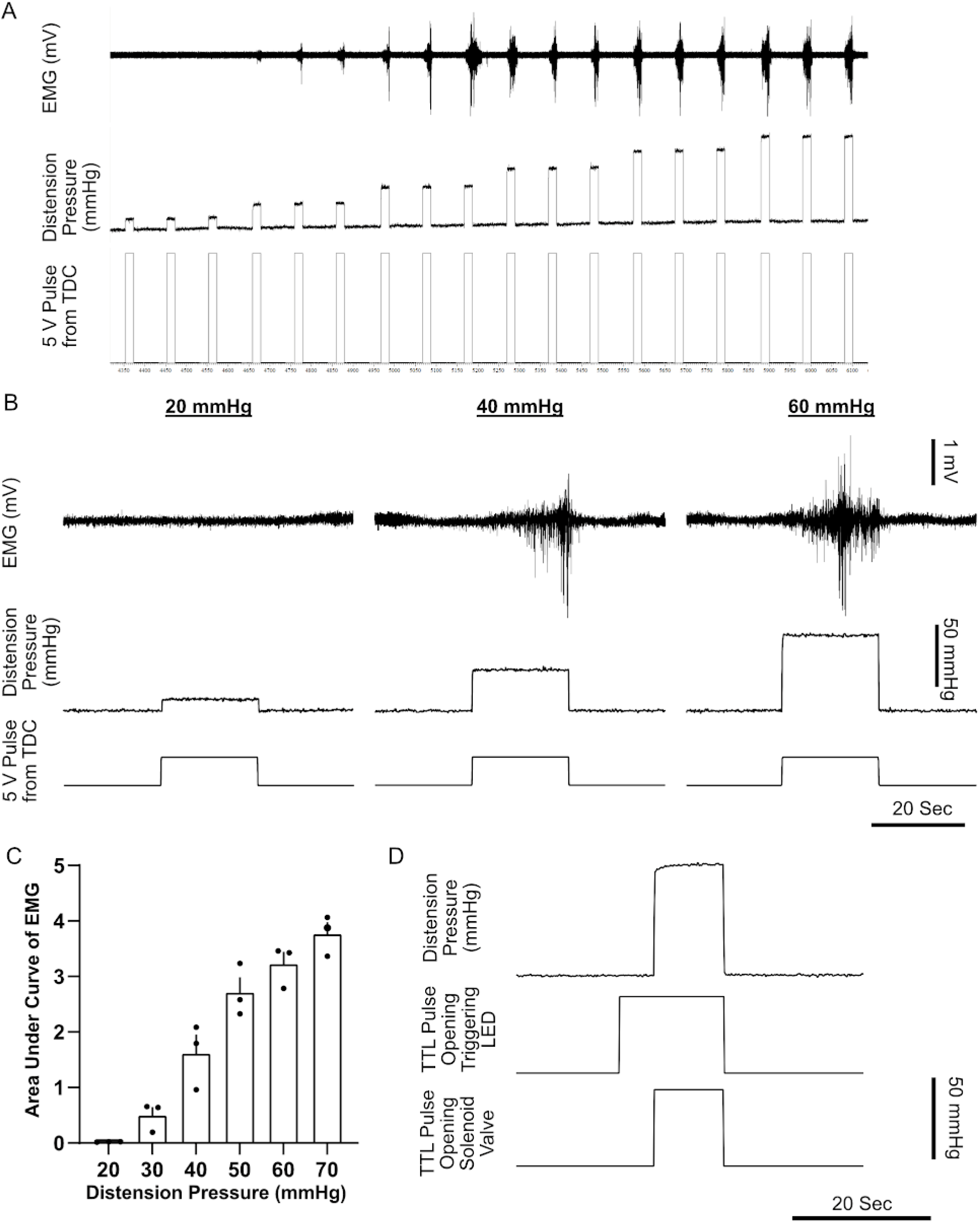
Validation of the Timed Distension Controller Using Mouse Bladder Distension to Evoke a Visceral Motor Response. **A**) An example screenshot of data acquisition software (Spike2, CED) used to acquire abdominal EMG responses evoked by bladder distension. The timed pressure controller is controlled using the user interface set 18 cycles of 20 sec off, 20 sec open, 60 seconds closed. The 5V pulse from the time pressure controller indicates the valve is open which correlates with the inline pressure transducer measurement (distension pressure) and physiologic EMG response. **B**) Representative examples of 20, 40 and 60 mmHg distensions demonstrating tight correlation between solenoid valve opening, don stream pressure and physiologic EMG response. **C**) Quantification of one mouse responses to all distensions. **D**) Example of TTL activation of the time distension controller and LED optogenetic stimulation using Spike2 and a Micro1401.

Our results indicate that the timed distension controller operates as designed, opening the solenoid at the indicated time and performing the input number of cycles (Figure 3a). The inline pressure transducer indicates the pressure increases at the same time the voltage pulse is sent. The distensions also correlate with increase physiologic EMG responses (Figure 3b and c), as has been demonstrated in numerous other publications [3, 11, 15]. These results indicate that the timed pressure delivery is working as intended, delivering air to distend the bladder and evoke abdominal EMG responses.

Next, we tested the TTL triggered mode of operation using Spike2 software to program the TTL pulses and a Micro 1404 to deliver the voltage stimulus to the timed pressure controller. As a proof in concept we also paired programed a TTL pulse to activate a blue laser prior to distension (Figure 3d). This can be used in optogenetic experiments to activate opsins, light activated proteins, in a population of cells prior to delivering the distension stimulus. Here we show a TTL pulse sent to the laser 5 seconds prior to sending the triggering stimulus to the timed distension controller. The timed distension controller opens the valve which correlates to in an increase in distension pressure (Figure 3d). When the valve closes (after 10 seconds) the distension quickly dissipates. Together these results show that by using the user interface we were able to control the solenoid valve and that delivery of bladder distension was able to replicate the capabilities of previous work.

## 8. Acknowledgements

We would like to acknowledge Alex Toney and Al Amin Hadis for their assistance with early hardware iterations. We would also like to thank Gabriella Robilotto for her technical support and logistical assistance for the project.

## Author contributions

ADM conceived of and designed the hardware; JH and NL designed and wrote the python software; TP wrote the Arduino software; TP designed the 3D printed case; TP and ADM, collected data, performed analysis, and wrote the manuscript.

## Data availability

All design files, and software are published and available at: https://osf.io/p6t2x/

## 10. Human and animal rights

All experiments were performed using adult (8-12 weeks old) female mice (B6/C57) housed in a facility on a 12-hour light/dark cycle with access to food and water ad libitum. All the procedures involving mice were approved by the University of Florida Institutional Animal Care and Use Committees, and in strict accordance with the US National Institute of Health (NIH) Guide for the Care and Use of Laboratory Animals (NIH Publications No. 8023, revised 1978).

